# Genome-wide identification and gene expression analysis of Clade A protein phosphatase 2C family genes in *Brassica juncea* var. *tumida*

**DOI:** 10.1101/2021.09.14.460405

**Authors:** Chunhong Cheng, Zhaoming Cai, Rongbin Su, Yuanmei Zhong, Li Chen, Lixia Wang, Guojian Chen, Lun Yan, Changman Li

## Abstract

Abscisic acid (ABA) plays crucial roles in plant response to environmental stresses and development. The clade A phosphatases (PP2Cs) play a crucial role in ABA signaling. However, little is known about the details regarding PP2Cs family genes in *Brassica juncea* var. *tumida*. Here, 20 clade A PP2Cs family genes were identified in tuber mustard genome, including *BjuABI1s*, *BjuABI2s*, *BjuAHG1s*, *BjuAHG3s*, *BjuHAB1*, *BjuHAB2s*, *BjuHAI1s*, *BjuHAI2s* and *BjuHAI3*. The promoters of BjuPP2Cs family genes contained various of cis-acting elements, such as ABRE, GT1GMSCAM4, ARFAT and MYB1AT. We also analyzed the expression pattern of clade A *BjuPP2Cs* under abiotic stresses (low temperature, NaCl and ABA) treatment, pathogen *Plasmodiophora brassicae* treatment and different stages of stem swollen. The results suggested that clade A *BjuPP2Cs* regulated tuber mustard response to *P. brassicae* to mediate the formation of clubroot and might play roles in stem swelling and response to abiotic stresses. This study provides valuable information for further functional investigations of clade A PP2Cs family genes in *B. juncea* var. *tumida*.

## Introduction

The reversible phosphorylation of proteins by protein phosphatases is a crucial process which modulates plant growth and development ^[1, 2]^. Protein phosphatases (PPs), removed the phosphate group of phosphorylated proteins, play important roles in regulation of protein function. PPs can be divided into two major subfamilies: protein tyrosine phosphatases and protein serine/threonine phosphatases. Based on the distinct amino acid sequences and crystal structures, the protein serine/threonine phosphatases can be classified into two groups: the phosphor-protein phosphatase (PPP) and the phosphoprotein metallophosphatases (PPM). The PP1, PP2A and PP2B subgroups belong to PPP family, whereas the PP2C subgroup belongs to the PPM family (Mg^2+^ or Mn^2+^ dependent) ^[3]^.

ABA plays a crucial role in plant response to environmental stresses such as abiotic stresses (salinity and low temperature stresses) and biotic stresses (such as pathogen) ^[4–6]^. Until now, the core components of the ABA signaling pathway include pyrabactin resistance (PYR)/PYR1-like (PYL) regulatory components of ABA receptor (RCAR) protein family (ABA receptors), the co-receptors clade A protein type 2C phosphatases (PP2Cs), and sucrose nonfermenting-1-related protein kinase class 2 (SnRK2s). In the absence of ABA, PP2Cs interact with SnRK2s to removing the phosphate group and inhibit the kinase activity of SnRK2s, which resulted in turning off ABA signaling pathway. In the presence of ABA, ABA binds the PYR/PYL/RCAR receptors and then interacts with PP2Cs, which inhibit the phosphatase activity of PP2Cs and eliminate the inhibitory effect of the phosphatases on SnRK2s to turn on the ABA signaling ^[7–9]^. In *Arabidopsis*, nine clade A PP2Cs (AtABI1, AtABI2, AtAHG1, AtHG3, AtHAB1, AtHAB2, AtHAI1, AtHAI2 and AtHAI3) have been identified as the negative regulators in ABA signaling pathway. The dominant mutations *abi1-1* and *abi2-1* showed the insensitive phenotype to ABA in germination and greening stages indicating that ABI1 and ABI2 were the negative regulators in ABA signaling ^[10]^. The *ahg1-1* and *ahg3-1* mutant showed ABA hypersensitivity phenotype in germination and post germination growth stages, but not in adult plants. And the expression level of *AHG1* was strongest in seeds indicating that it played an important role in response to ABA in seed ^[11, 12]^. *HAB1* underwent alternative splicing and produced two splice variants, named *HAB1.1* and *HAB1.2*. HAB1.1 and HAB1.2 played the opposing roles in ABA-mediated seed germination and post-germination stages ^[13]^. In *B.rapa* the PP2C genes were structurally conserved based on the amino acid sequence alignment, phylogenetic analysis and conserved domains; and the gene expression levels were induced by heat, cold, ABA and drought treatment ^[14]^. Twenty group A PP2C homologous genes of *B. oleracea* were identified; the genetic analysis corroborated the presence of two to three gene copies in *B. oleracea* in comparison to clade A PP2C genes in *Arabidopsis thaliana*; the gene expression patterns of *PP2Cs* in *B. oleracea* were significant differences ^[15]^.

Tuber mustard, *Brassica juncea* var. *tumida* (AABB, 2n = 36), an allotetraploid species, which belongs to *Brassicaceae*, came from a natural cross between *B. rapa* (AA, 2n = 20) and *B. nigra* (BB, 2n = 16), followed by chromosome doubling ^[16]^. The swollen stem of tuber mustard is the raw material of Fuling mustard and is an important vegetable. However, little is known about the regulation mechanism of stem swelling ^[17]^. And during the growth and development stages, tuber mustard frequently suffers from abiotic and biotic stresses, such as salt stress, cold stress and *Plasmodiophora brassicae*, which leads to the suppression of plant growth and development and the formation of clubroot. Clubroot is one of the most serious biotic stresses that plants need to cope with and usually results in the suppression of tuber mustard growth and stem swollen, leading to limitation in yield ^[18]^. Therefore, illustrating the mechanisms underlying the stem swollen and the resistance to abiotic and biotic stresses will be helpful for improving the production of tuber mustard. Clade A PP2Cs of ABA signaling pathway play important roles in response to abiotic and biotic stresses and regulation plant development. However, the gene structures, protein motifs, gene duplication events and functions of clade A *PP2Cs* in *B. juncea* var. *tumida* remains mainly unknown.

In this study, twenty clade A PP2C family genes were identified in *B. juncea* var. *tumida* genome. The phylogenic relationship, gene structures, and protein motifs were compared between clade A *BjuPP2Cs* and *AtPP2Cs* and found that they shared similar gene characteristics. Following the analysis of gene features, we analyzed the transcript levels of clade A *BjuPP2Cs* under *P. Brassicae*, NaCl, low temperature and ABA treatment and different stages of stem swollen. The results showed that clade A *BjuPP2Cs* were induced by pathogen and abiotic stresses and induced in the stages of stem swollen, suggesting that clade A BjuPP2C family genes played crucial roles in plant response to abiotic stresses, pathogen *P. Brassicae* and regulation of stem swollen.

## Materials and Methods

### Materials and growth conditions

The tuber mustard cultivar Yong’an was used in this study. The seeds were sterilized and sowed in MS medium. The growth room was at 22°C and 6000 lx (long-day conditions; 16 h light/8 h dark). To analyze the gene expression patterns of clade A BjuPP2Cs genes under biotic stress, 14-day-old seedlings were treated with *P. brassicae* liquid (OD_600_=0.07) for 0, 0.25, 0.5, 1, 3, 5, 7, and 9 days. To analysis the gene expression level in different growth stages of stem swelling, the samples D1 (the stems of 1-month-old seedlings, six leaf stage), D2 (the stems of 2-month-old seedlings, primary stage of stem swelling), D3 (the stems of 3-month-old seedlings, early stage of stem swelling), D4 (the stems of 4.5-month-old seedlings, fast-growing stage of stem swelling) and D5 (the stems of 5-month-old seedlings, last stage of stem swelling) of *B. juncea* var. *tumida* which were grown in the field were collected. To analyze the gene expression patterns of clade A BjuPP2Cs genes under abiotic stresses, 5-day-old seedlings were treated with 50 μ M ABA, 100 mM NaCl and 4°C treatment for 3 h.

### Bioinformatics analysis

The gene sequences of clade A *AtPP2Cs* and their homologous genes in tuber mustard were searched in TAIR (http://www.arabidopsis.org/) and *Brassica Databases* (http://brassicadb.cn/#/). The phylogenic tree was analysis with the neighbor-joining method by MEGA5. Gene structures were analyzed by Gene Structure Display Server 2.0 (http://gsds.cbi.pku.edu.cn/). The promoter regions of clade A *BjuPP2Cs* were chose by Flanking region in *Brassica Database*, and the 2 kb regions upstream of ATG contained no other genes. So the regions located 2 kb upstream of the clade A *BjuPP2Cs* coding sequences were used as the promoter sequences, and promoter cis-element analysis was performed using New PLACE (https://www.dna.affrc.go.jp/PLACE/?action=newplace). Protein domain was analysis by SMART (http://smart.embl-heidelberg.de/), ExPASy (http://prosite.expasy.org/prosite.html) and ESPrint3.0 (https://espript.ibcp.fr/ESPript/cgi-bin/ESPript.cgi).

### Gene expression analysis

Total RNA of tuber mustard was extracted under pathogen and abiotic stresses treatment. qRT-PCR was performed using the cDNA. The transcript level was analysis by the comparative CT method, and *BjuActin3* was used as the internal reference. The qRT-PCR experiments were carried out three times with three replicates each. The primers used in this study were listed in Table S1.

### Statistical analysis

All data were analyzed using SigmaPlot 10.0 (Systat Software, Inc., Chicago, IL) and SPSS 16.0 software. The averages and standard deviations of all results were calculated, and for multiple groups of samples, the one-way ANOVA followed by the Dunnett test was used. The statically significant treatments were marked with ‘***’ (P<0.001), ‘**’ (0.001<P<0.01) and ‘*’ (0.01<P<0.05).

## Results

### Genome-wide identification and characterization of clade A *BjuPP2Cs* in *B. juncea* var. *tumida*

Twenty genes as homologs of clade A *PP2Cs* were identified in *B. juncea* var. *tumida* through BLASTP in *Brassica* database using nine clade A AtPP2C protein sequences as references, and the naming rules based on the names of PP2Cs in *Arabidopsis thaliana*, such as the five copies of AtABI1 named BjuABI1-1, BjuABI1-2, BjuABI1-3, BjuABI1-4 and BjuABI1-5 (Table 1). The gene lengths ranged from 1288 bp to 5344 bp with 2-8 exons in each sequence. The protein lengths of these twenty PP2C homologs ranged from 325 (BjuABI1-4) to 606 (BjuAHG3-3) amino acid (aa) residues. The relative molecular weights of those proteins varied from 35.907 kD (BjuABI1-4) to 67.615 kD (BjuAHG3-3), and the isoelectric point (PI) ranged from 4.40 to 7.51 (Table 1). The twenty *BjuPP2C* genes were distributed in 13 of the 18 chromosomes of *B. juncea* var. *tumida*. Each of the chromosomes A01, A05, A08, A09, B01, B05, B06, B07, and B08 contained one gene, each of the chromosomes A03 and B02 contained two genes, and each of the chromosomes A10 and B03 contained three genes (Figure 1).

**Figure 1.**
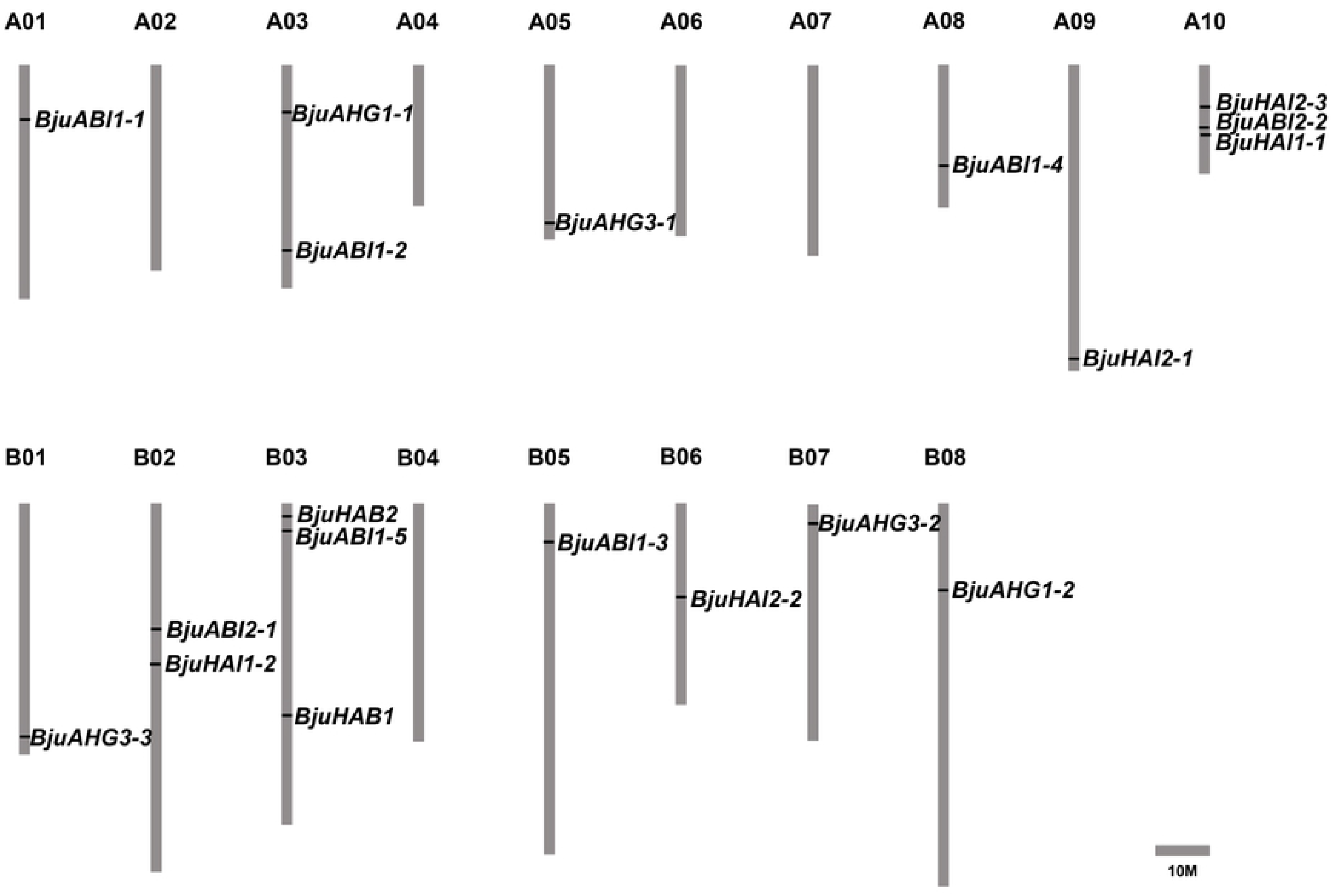
The distribution of clade A *BjuPP2Cs* in *B. juncea* var. *tumida* chromosomes. Twenty identified clade A *BjuPP2Cs* were mapped to the 13 of 18 chromosomes. The chromosome name is at the top of each bar. The scale of the chromosome is in millions of bases (Mb).

**Table 1.**
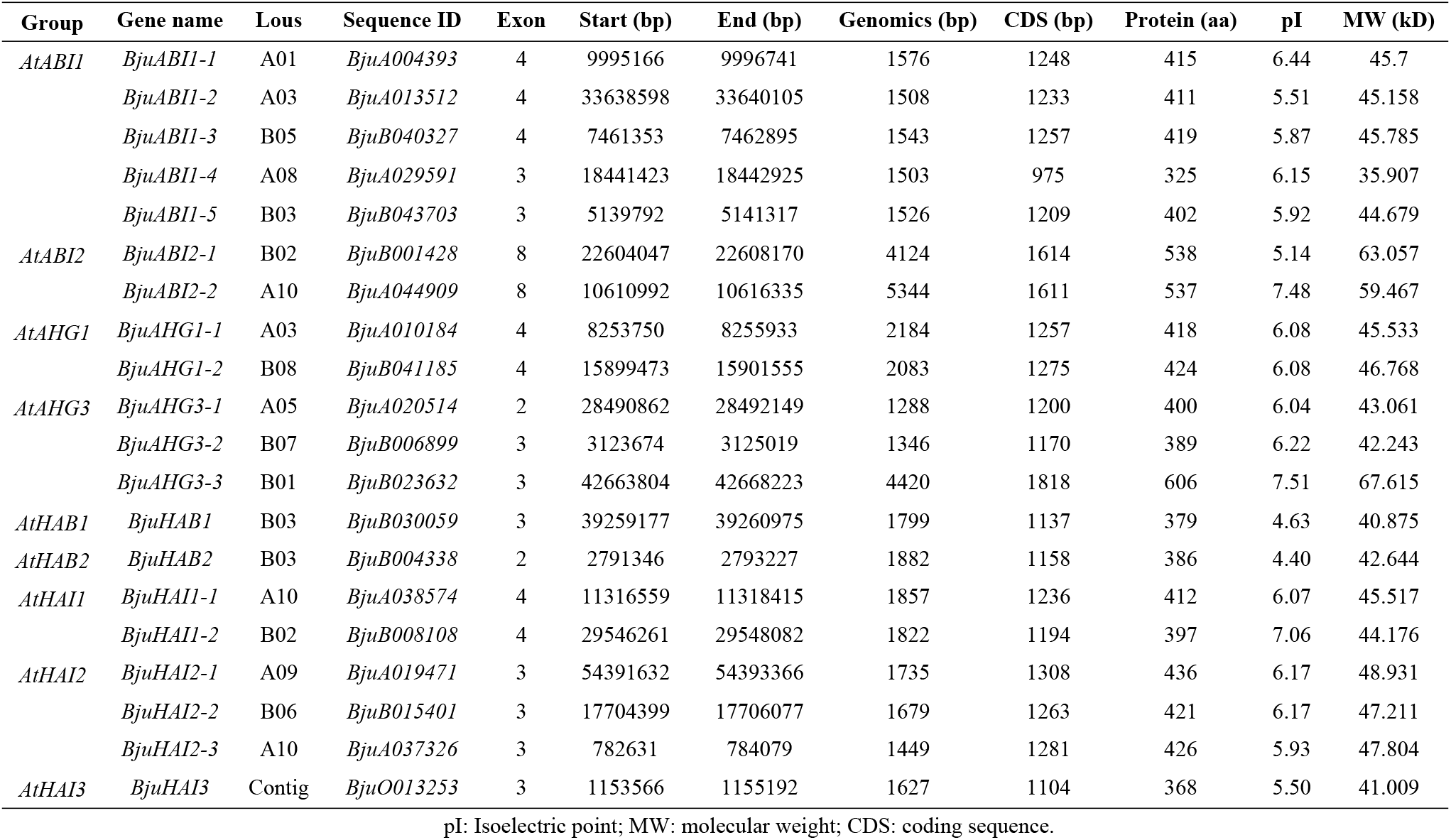
The clade A BjuPP2Cs family members in *B. juncea* var. *tumida*.

| Group | Gene name | Lous | Sequence ID | Exon | Start (bp) | End (bp) | Genomics (bp) | CDS (bp) | Protein (aa) | pI | MW (kD) |
| --- | --- | --- | --- | --- | --- | --- | --- | --- | --- | --- | --- |
| *AtABI1* | *BjuABI1-1* | A01 | *BjuA004393* | 4 | 9995166 | 9996741 | 1576 | 1248 | 415 | 6.44 | 45.7 |
|  | *BjuABI1-2* | A03 | *BjuA013512* | 4 | 33638598 | 33640105 | 1508 | 1233 | 411 | 5.51 | 45.158 |
|  | *BjuABI1-3* | B05 | *BjuB040327* | 4 | 7461353 | 7462895 | 1543 | 1257 | 419 | 5.87 | 45.785 |
|  | *BjuABI1-4* | A08 | *BjuA029591* | 3 | 18441423 | 18442925 | 1503 | 975 | 325 | 6.15 | 35.907 |
|  | *BjuABI1-5* | B03 | *BjuB043703* | 3 | 5139792 | 5141317 | 1526 | 1209 | 402 | 5.92 | 44.679 |
| *AtABI2* | *BjuABI2-1* | B02 | *BjuB001428* | 8 | 22604047 | 22608170 | 4124 | 1614 | 538 | 5.14 | 63.057 |
|  | *BjuABI2-2* | A10 | *BjuA044909* | 8 | 10610992 | 10616335 | 5344 | 1611 | 537 | 7.48 | 59.467 |
| *AtAHG1* | *BjuAHG1-1* | A03 | *BjuA010184* | 4 | 8253750 | 8255933 | 2184 | 1257 | 418 | 6.08 | 45.533 |
|  | *BjuAHG1-2* | B08 | *BjuB041185* | 4 | 15899473 | 15901555 | 2083 | 1275 | 424 | 6.08 | 46.768 |
| *AtAHG3* | *BjuAHG3-1* | A05 | *BjuA020514* | 2 | 28490862 | 28492149 | 1288 | 1200 | 400 | 6.04 | 43.061 |
|  | *BjuAHG3-2* | B07 | *BjuB006899* | 3 | 3123674 | 3125019 | 1346 | 1170 | 389 | 6.22 | 42.243 |
|  | *BjuAHG3-3* | B01 | *BjuB023632* | 3 | 42663804 | 42668223 | 4420 | 1818 | 606 | 7.51 | 67.615 |
| *AtHAB1* | *BjuHAB1* | B03 | *BjuB030059* | 3 | 39259177 | 39260975 | 1799 | 1137 | 379 | 4.63 | 40.875 |
| *AtHAB2* | *BjuHAB2* | B03 | *BjuB004338* | 2 | 2791346 | 2793227 | 1882 | 1158 | 386 | 4.40 | 42.644 |
| *AtHAI1* | *BjuHAI1-1* | A10 | *BjuA038574* | 4 | 11316559 | 11318415 | 1857 | 1236 | 412 | 6.07 | 45.517 |
|  | *BjuHAI1-2* | B02 | *BjuB008108* | 4 | 29546261 | 29548082 | 1822 | 1194 | 397 | 7.06 | 44.176 |
| *AtHAI2* | *BjuHAI2-1* | A09 | *BjuA019471* | 3 | 54391632 | 54393366 | 1735 | 1308 | 436 | 6.17 | 48.931 |
|  | *BjuHAI2-2* | B06 | *BjuB015401* | 3 | 17704399 | 17706077 | 1679 | 1263 | 421 | 6.17 | 47.211 |
|  | *BjuHAI2-3* | A10 | *BjuA037326* | 3 | 782631 | 784079 | 1449 | 1281 | 426 | 5.93 | 47.804 |
| *AtHAI3* | *BjuHAI3* | Contig | *BjuO013253* | 3 | 1153566 | 1155192 | 1627 | 1104 | 368 | 5.50 | 41.009 |

### Phylogenic analysis and gene structures of clade A PP2C family genes

To analyze the evolutionary relationships between PP2C homologs in *B. juncea* var. *tumida* and *Arabidopsis thaliana*, a phylogenetic tree was constructed by MEGA5 software using the neighbor-joining method. According to the phylogenic tree, twenty clade A BjuPP2Cs with nine AtPP2Cs were identified and clustered into nine clades. The first clade was AtHAI2 and three homologs BjuHAI2-1, BjuHAI2-2 and BjuHAI2-3; the second clade was AtHAI3 and BjuHAI3; the third clade was AtHAI1 and its two homologs BjuHAI1-1 and BjuHAI1-2; AtAHG3 and its homologs BjuAHG3-1, BjuAHG3-2 and BjuAHG3-3 were clustered into one clade; AtAHG1, BjuAHG1-1 and BjuAHG1-2 were clustered into one clade; AtHAB1 and BjuHAB1 were clustered into one clade; AtHAB2 and BjuHAB2 were clustered into one clade; the eighth clade was AtABI2 and its two homologs BjuABI2-1 and BjuABI2-2; and the last clade was AtABI1 and its homologs BjuABI1-1, BjuABI1-2, BjuABI1-3, BjuABI1-4 and BjuABI1-5 (Figure 2). Clade A PP2C family genes in the same subfamilies may have similar functions. To understand their gene structures, we analyzed the gene exon-introns. According to the results, *AtHAI1* and *BjuHAI1s*, *AtHAI2* and *BjuHAI2s*, *AtHAI3* and *BjuHAI3s*, *AtAHG1* and *BjuAHG1s* had the same gene structures, respectively; *AtAHG3* contained 4 exons, while *BjuAHG3-1, BjuAHG3-2* and *BjuAHG3-3* had 2, 3 and 3 exons, respectively; *AtHAB1* and *BjuHAB1* contained 4 and 3 exons, respectively; *AtHAB2* and *BjuHAB2* contained 4 and 2 exons, respectively; *BjuABI2-1* and *BjuABI2-2* all contained 8 exons with the exception of *AtABI2* (4 exons); *AtABI1*, *BjuABI1-1*, *BjuABI1-2* and *BjuABI1-3* all had 4 exons, in contrast, *BjuABI1-4* and *BjuABI1-5* contained 3 exons (Figure 2).

**Figure 2.**
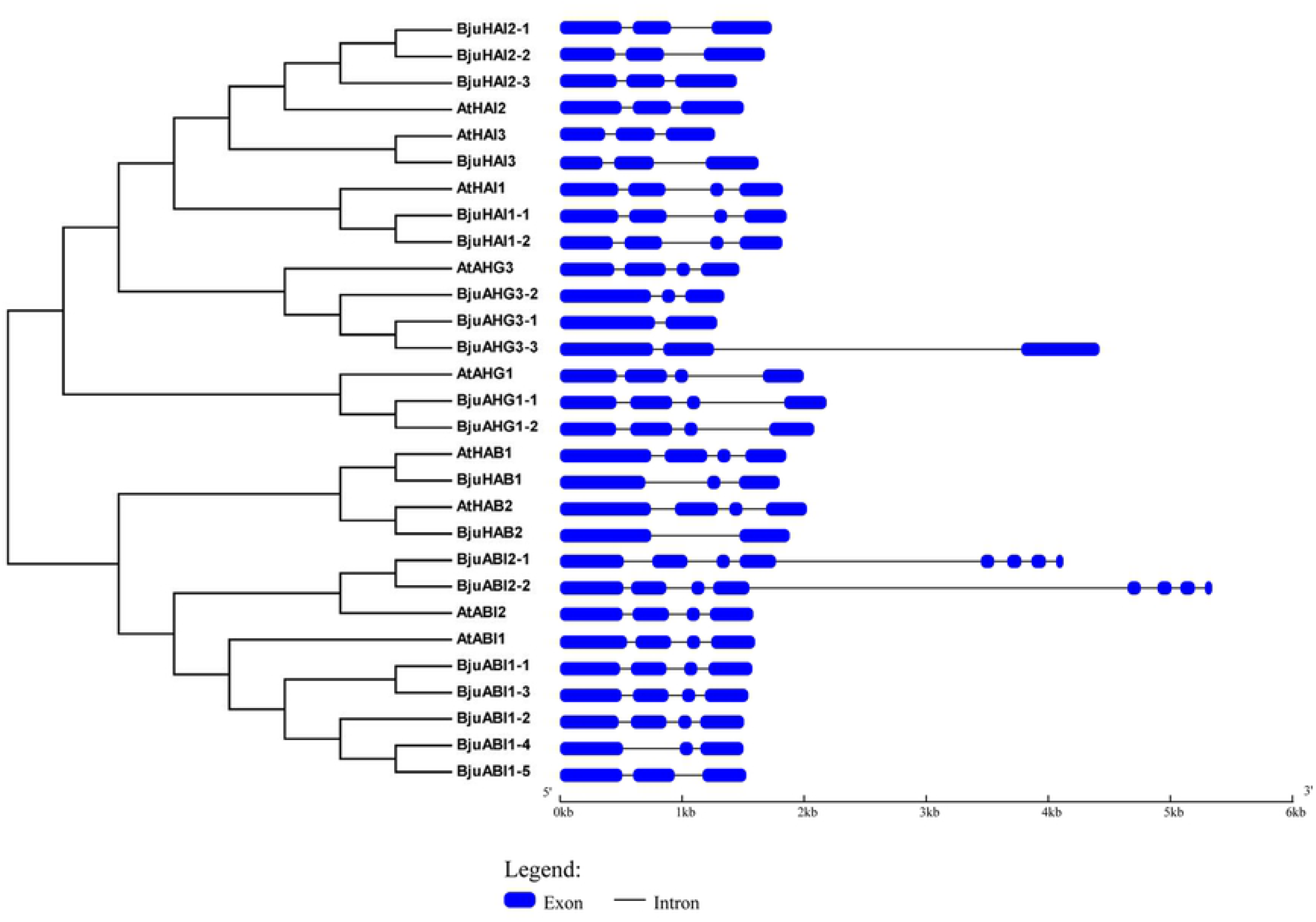
The phylogenic tree and gene structures of clade A *PP2C* family genes. The phylogenic tree was built using the neighbor-joining (NJ) method and the exon-intron structure of clade A *PP2C* homologs was drawn according to their phylogenic relationships. The blue boxes and black lines denoted exons and introns, respectively.

### The conserved motif analysis of clade A PP2C homologs

To analysis the protein function of PP2C homologs, protein conserved motifs were conducted. The results showed that all the clade A BjuPP2Cs contained PP2Cc domain (Serine/threonine phosphatases, family2C, catalytic domain), which was consistent with clade A AtPP2Cs; only BjuABI2-1 and BjuABI2-2 contained transmembrane domain (Figure 3). As some clade A *BjuPP2Cs* lacked of some exons, such as *BjuABI1s* and *BjuAHG3s*, might lead to the different effects on the catalytic PP2C domain in their proteins. So we analyzed the motif sequences of BjuPP2Cs by ClustalX and ESPrint 3.0 (https://espript.ibcp.fr/ESPript/cgi-bin/ESPript.cgi). The protein serine/threonine phosphatase 2C (PP2C) family contained 11 conserved motif, named motif 1-5, motif 5a, motif 5b and motif 6-11 ^[19]^. In our result, all of the 11 conserved motifs in the clade A BjuPP2C proteins were identified, suggesting that the catalytic PP2C domain in their proteins were conserved even if they had different gene structures (Fig. S1).

**Figure 3.**
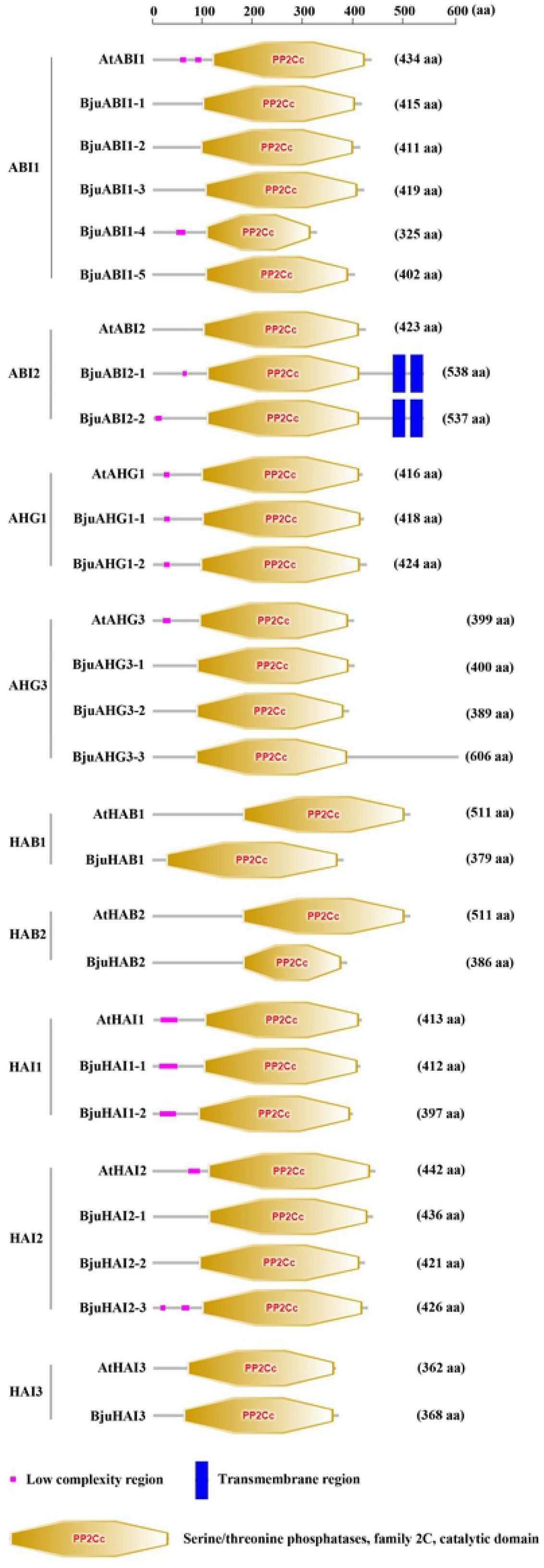
The conserved motifs of clade A PP2C homologs proteins. These motifs were identified using the online software of SMART, and colored boxes indicated conserved motifs and black lines represent non-conserved sequences.

### Promoter cis-acting regulatory elements prediction of clade A BjuPP2C homologs

To further understand the potential roles of clade A *BjuPP2Cs*, the promoters of clade A *BjuPP2Cs* were analyzed. The promoter regions of clade A *BjuPP2Cs* were chose by Flanking region in Brassica Database, and the 2 kb regions upstream of ATG contained no other genes. So we chose the 2000 bp DNA fragment upstream of the ATG start codon as the promoter sequences and analyzed promoter cis-elements. The results showed that all *BjuPP2Cs* contained at least one hormone-related elements in the promoters such as ARFAT (TGTCTC, responsive to auxin) ^[20]^ and ASF1MOTIFCAMV (TGACG, responsive to auxin and salicylic acid) ^[21]^ (Figure 4). The promoters of all *BjuPP2Cs* genes, except *BjuHAI1-2*, contained 1 to 7 cis-acting elements related to ABA, such as ABRE (ACGTG, responsive to Abscisic acid, ABA) ^[22]^ (Figure 4). In addition, the promoters of the BjuPP2Cs homologs all contained stressed-related elements, such as the MYCCONSENSUSAT (CANNTG, responsive to dehydration stress) ^[23]^, MYBATRD22 (CTAACCA, responsive to dehydration stress) ^[24]^, MYB1AT (WAACCA, responsive to dehydration stress) ^[23]^, CBFHV (RYCGAC, responsive to dehydration stress) ^[25]^, CRTDREHVCBF2 (GTCGAC, responsive to low temperature) ^[26]^, LTRECOREATCOR15 (CCGAC, responsive to low temperature) ^[27]^, GT1GMSCAM4 motif (GAAAAA, responsive to pathogen and salt stress) ^[28]^, GCCCORE (GCCGCC, responsive to pathogen) ^[29]^ and MYB1LEPR (GTTAGTT, responsive to defence) ^[30]^ (Figure 4). Together, the promoters of BjuPP2Cs homologs contained diverse cis-elements responsive to ABA, auxin, dehydration, low temperature and pathogen, indicating that the genes expression of clade A BjuPP2Cs homologous were regulated by hormone, biotic stresses and abiotic stresses, and clade A *BjuPP2Cs* might play a role in regulating tuber mustard response to hormone and stresses.

**Figure 4.**
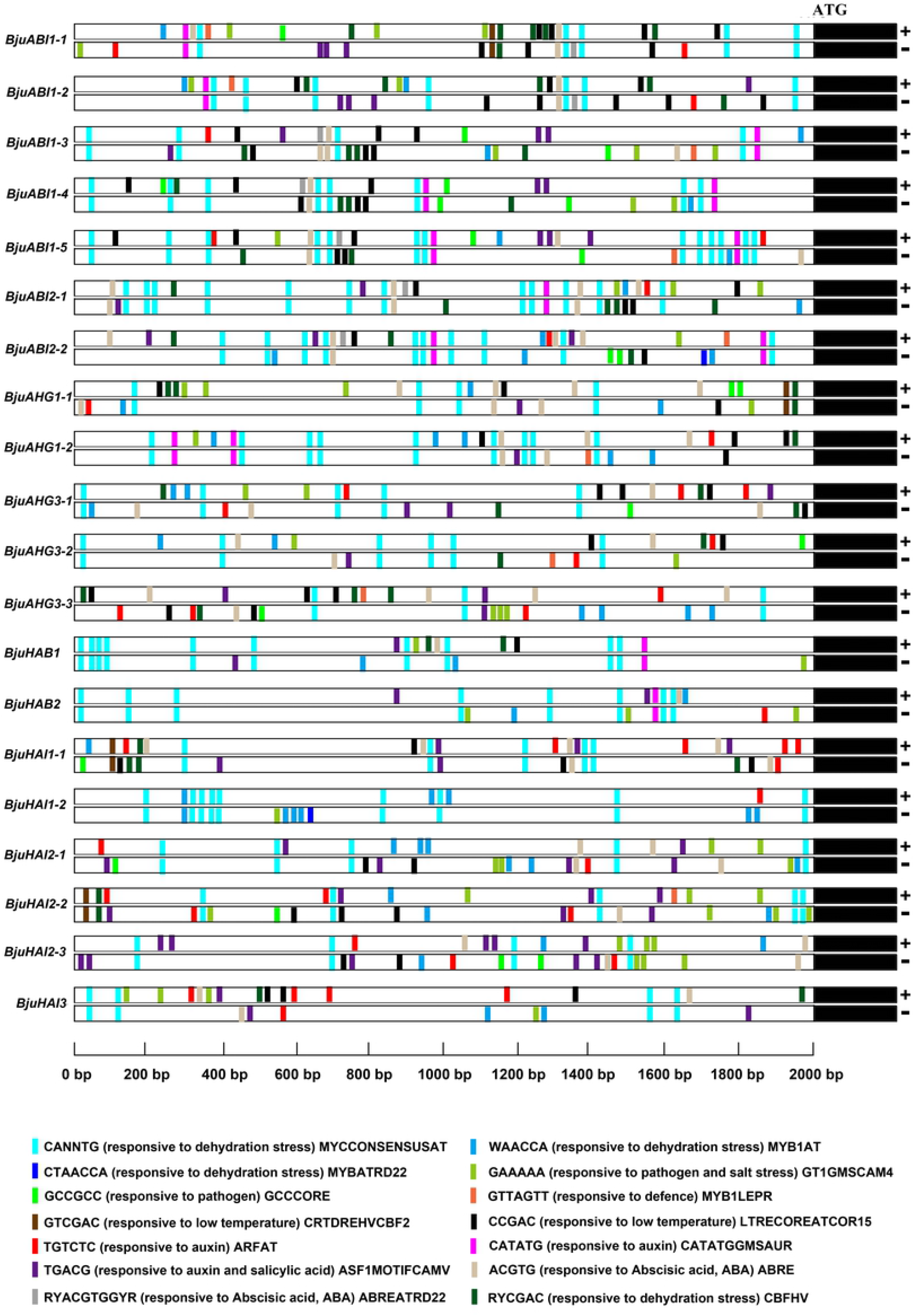
The promoter *cis*-elements analysis of clade A *BjuPP2Cs*. The 2 kb DNA fragments upstream of the ATG staring code of clade A *BjuPP2Cs* were analyzed using online analysis software PlantCARE and PLACE.

### Tissue specific expression pattern analysis of clade A *BjuPP2Cs*

To investigate the tissue specific expression patterns of clade A *BjuPP2Cs*, we analyzed the gene expression levels at different growth stages and tissues (root, stem, swollen stem, leaf, flower and pod) using qRT-PCR. According to the results, clade A *BjuPP2Cs* were expressed in multiple tissues. Interestingly, the expression levels of *BjuABI1-1*, *BjuABI1-5*, *BjuABI2-1*, *BjuABI2-2*, *BjuAHG1-1*, *BjuAHG3-3*, *BjuHAI1-1*, *BjuHAI2-1*, *BjuHAI2-3* and *BjuHAI3* showed significant difference between stem and swollen stem, indicating that these clade A *BjuPP2Cs* might play a role in the regulation of stem swollen (Figure 5). The expression levels of *BjuABI1-2*, *BjuABI1-3*, *BjuABI1-4*, *BjuAHG3-3*, *BjuHAB1*, *BjuHAB2*, *BjuHAI1-1* and *BjuHAI2-1* were higher in root than that in other organs. Almost all the clade A BjuPP2Cs were expressed very low in flower and pod, especially *BjuHAI2-1* and *BjuHAI2-2* (nearly no expression) (Figure 5). The expression levels of the clade A *BjuPP2Cs* varied in different tissues and organs, indicating that they might play different roles in different organs. In addition, the different expression patterns of the same gene in different tissues and organs suggested that the expression patterns of the genes were existence of space-time specificity.

**Figure 5.**
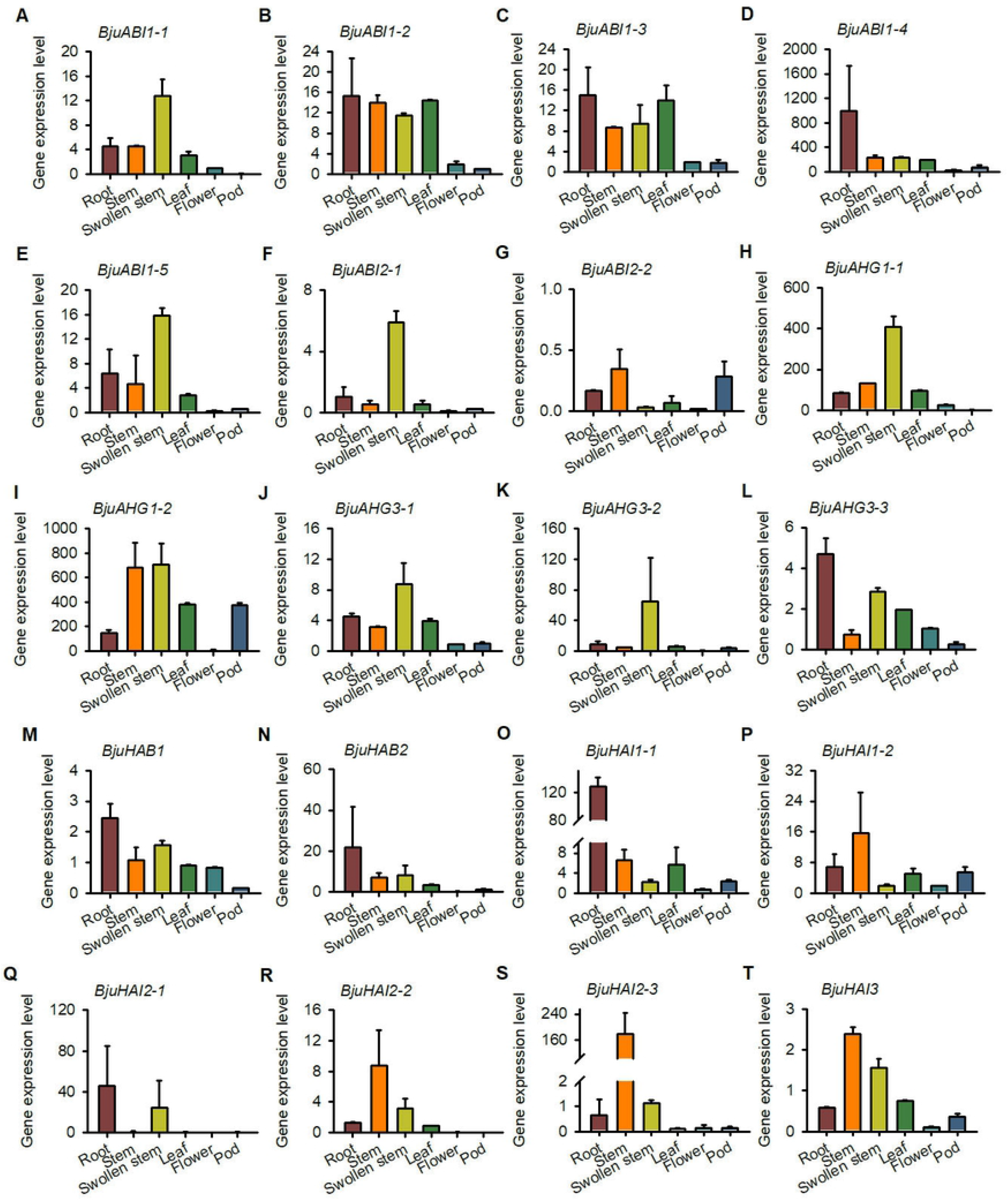
Expression patterns of clade A *BjuPP2Cs* in different tissues. Tissue specific expression pattern of clade A *BjuPP2Cs* were analyzed by qPCR. Data were normalized to the expression level of *BjuActin3*. The values are means ± standard error. Three independent biological repeats were performed.

### The expression patterns of clade A *BjuPP2Cs* in *B. juncea* var. *tumida* under abiotic stresses

To further analysis the expression levels of clade A *BjuPP2Cs* under abiotic stresses treatment, qRT-PCR assays were performed using 5-day-old tuber mustard seedlings treated with 50 μ M ABA, 100 mM NaCl and 4 °C treatment for 3 h. The qRT-PCR results showed that all of the clade A *BjuPP2Cs* genes were induced by NaCl significantly (Figure 6). Under low temperature treatment, the expression levels of *BjuABI1-2* to *BjuABI1-4*, *BjuABI2-2*, *BjuAHG1-2*, *BjuHAB2*, *BjuHAI2-3* and *BjuHAI3* were induced, whereas, the expression levels of *BjuABI2-1, BjuAHG3-1* to *BjuAHG3-3* and *BjuHAI2-1* were suppressed (Figure 6). Under ABA treatment, *BjuABI1-1* to *BjuABI1-4*, *BjuABI2-2*, *BjuAHG1-2*, *BjuAHG3-2*, *BjuAHG3-3*, *BjuHAB1*, *BjuHAI1-2* and *BjuHAI2-3* were induced by ABA treatment, however, *BjuABI2-1, BjuAHG3-1* and *BjuHAB2* were inhibited (Figure 6).

**Figure 6.**
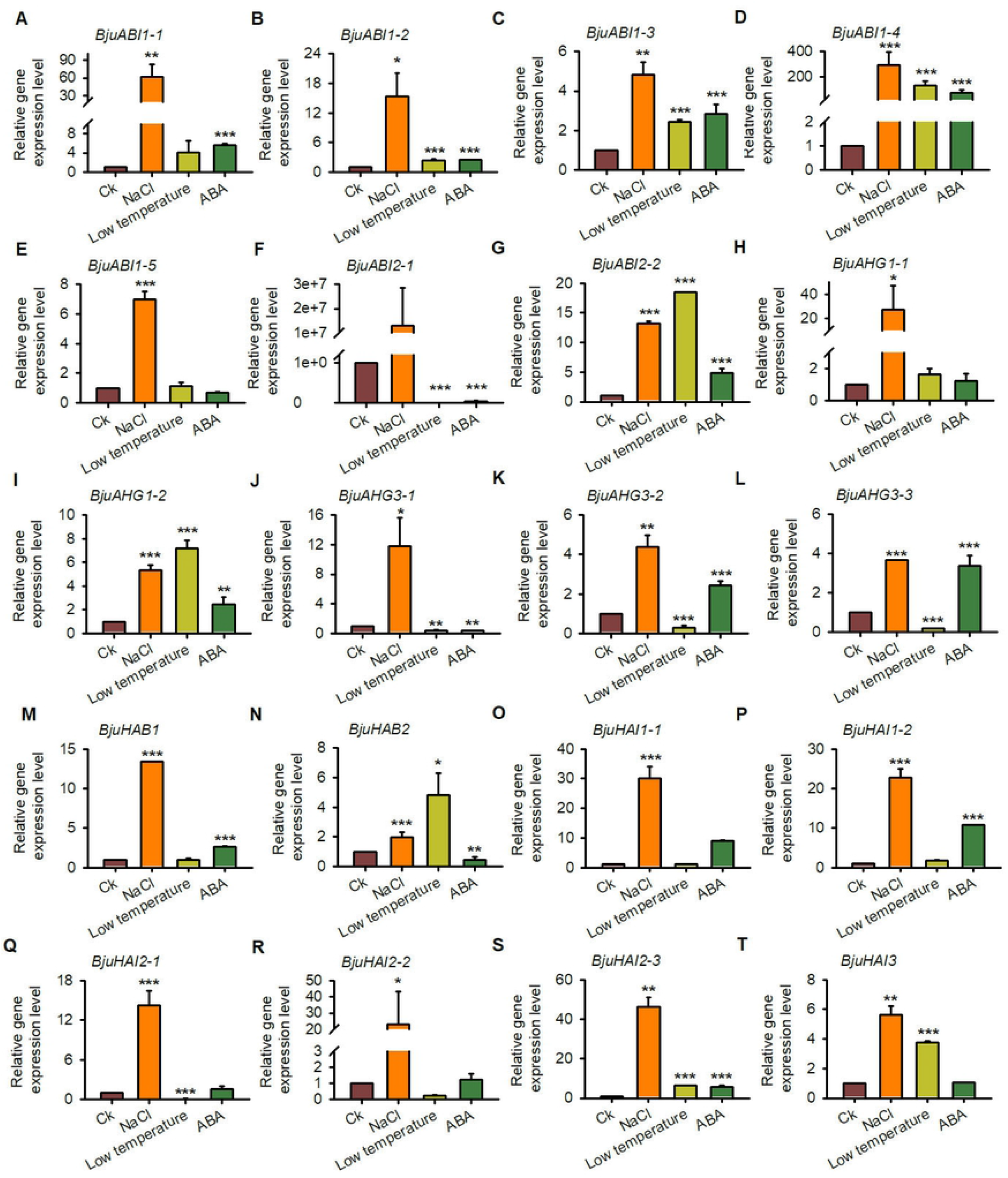
The expression patterns of clade A *BjuPP2Cs* under abiotic stresses. Total RNA was extracted from tuber mustard seedlings treated with ABA, NaCl and 4 °C for 3 h. Data were normalized to the expression level of *BjuActin3*. The values are means ± standard error. Three independent biological repeats were performed.

### The expression patterns of clade A *BjuPP2Cs* in *B. juncea* var. *tumida* under *P. brassicae* stress

To further analysis the gene expression patterns of clade A *BjuPP2Cs* in tuber mustard under pathogen stress, qRT-PCR assays were performed using tuber mustard seedlings treated with *P. brassicae*. Under *P. brassicae* treatment, 2-week-old tuber mustard seedlings were treated with *P. brassicae* for 0, 0.25, 0.5, 1, 3, 5, 7 and 9 days, and the qRT-PCR results showed that *BjuABI1-5*, *BjuAHG1-1*, *BjuAHG3-2*, *BjuAHG3-3*, *BjuHAB1*, *BjuHAB2*, *BjuHAI1-1*, *BjuHAI2-1*, *BjuHAI2-3* and *BjuHAI3* were highly induced by pathogen, especially on day 0.5 after pathogen treatment. The result indicated that *BjuABI1-5*, *BjuAHG1-1*, *BjuAHG3-2*, *BjuAHG3-3*, *BjuHAB1*, *BjuHAB2*, *BjuHAI1-1*, *BjuHAI2-1*, *BjuHAI2-3* and *BjuHAI3* may play roles in *P. brassicae* tolerance (Figure 7).

**Figure 7.**
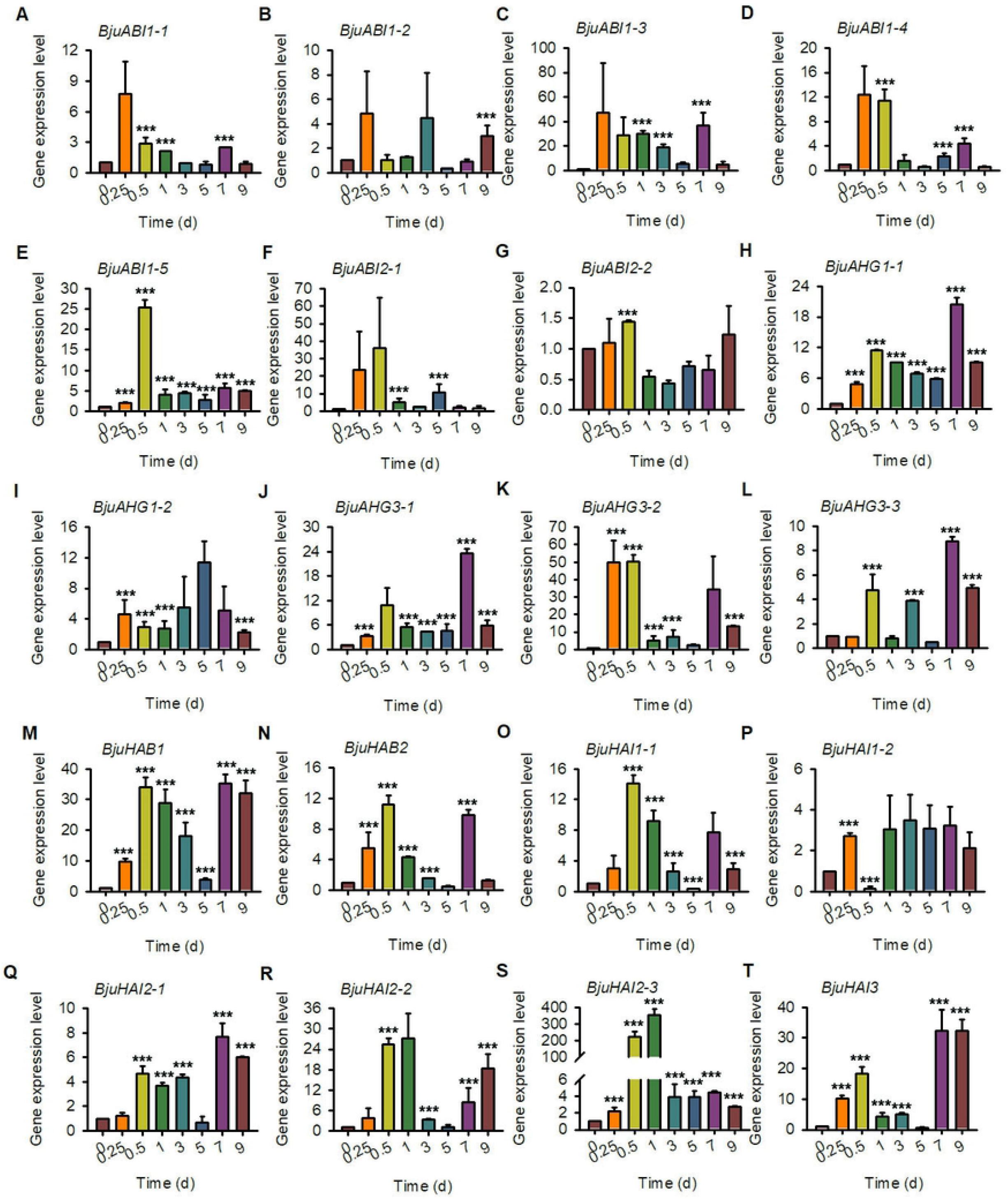
The expression patterns of clade A *BjuPP2Cs* under pathogen treatment. Total RNA was extracted from tuber mustard seedlings treated with *P. Brassicae* at the indicated time points. Data were normalized to the expression level of *BjuActin3*. The values are means ± standard error. Three independent biological repeats were performed.

### Gene expression levels of clade A *BjuPP2Cs* in *B. juncea* var. *tumida* during stem swelling stages

The swollen stem is the raw material of “Fuling Mustard”, however, the regulation mechanism of stem swelling was still undiscovered. To further explore the roles of clade A *BjuPP2Cs* in regulating the formation and development of swollen stem in tuber mustard, we collected the stems of *B. juncea var. tumida*, which were taken in the field at different growth and development stages. The stems were named as D1 (the stems of 1 month old seedlings, six leaf stage), D2 (the stems of 2 month old seedlings, primary stage of stem swelling), D3 (the stems of 3 month old seedlings, early stage of stem swelling), D4 (the stems of 4.5 month old seedlings, fast growing stage of stem swelling), and D5 (the stems of 5 month old seedlings, last stage of stem swelling). The qRT-PCR assay was performed and the results showed that the gene expression levels of *BjuABI1-1*, *BjuAHG1-1*, *BjuHAI2-2*, *BjuHAI2-3* and *BjuHAI3* were inhibited during stem swelling stages (D2-D4 stages). The expression levels of *BjuABI1-3*, *BjuABI2-1*, *BjuHAB1* and *BjuHAI1-1* were induced during stem swelling stages (D2-D4 stages) (Figure 8). The results suggested that clade A *BjuPP2Cs* may play roles in the regulation of the formation and development of swollen stem.

**Figure 8.**
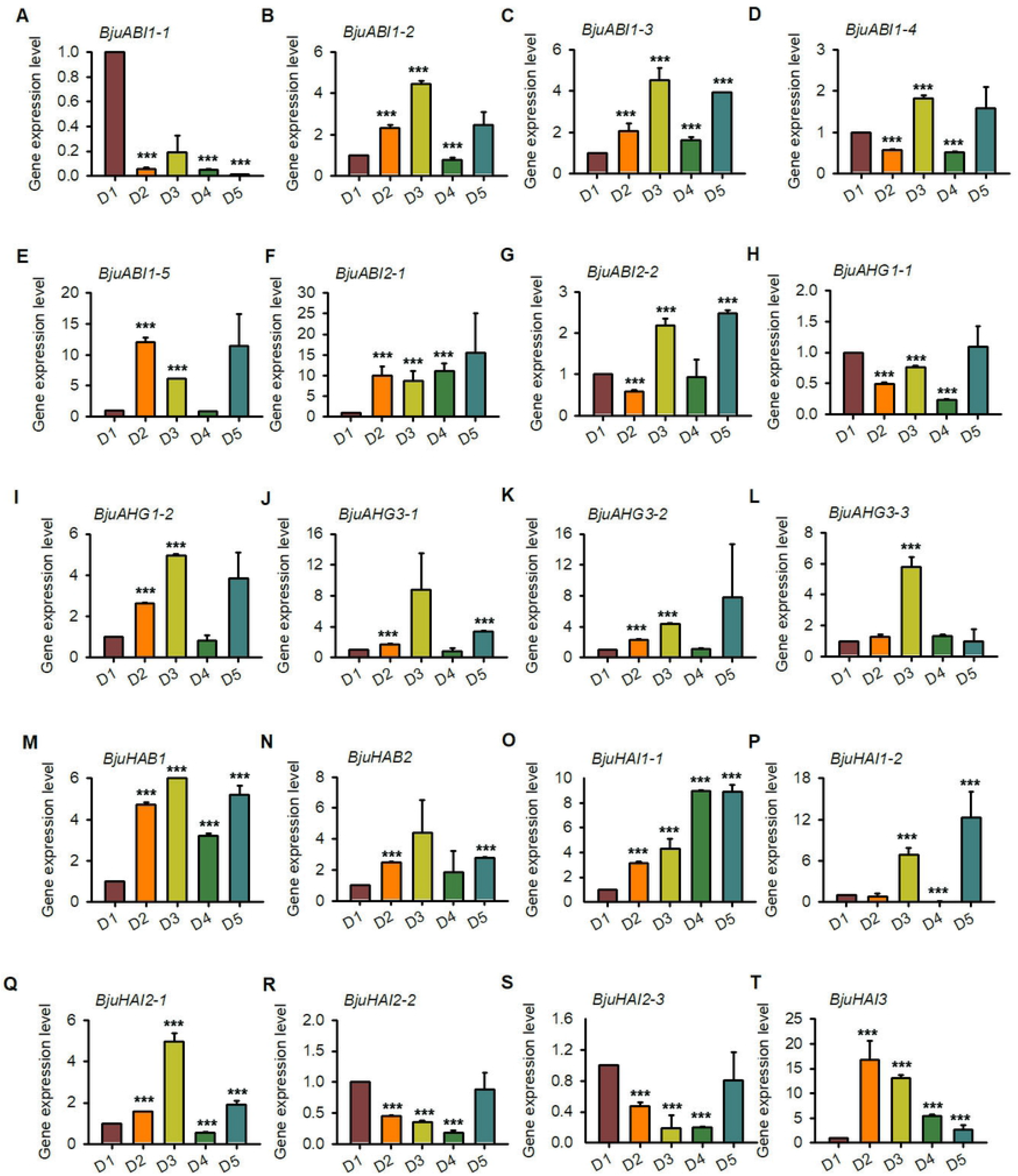
The expression patterns of clade A *BjuPP2Cs* at different stages of stem swelling. Total RNA was extracted from the seedlings of D1, D2, D3, D4 and D5 stages. Data were normalized to the expression level of *BjuActin3*. The values are means ± standard error. Three independent biological repeats were performed.

## Discussion

The swollen stem of *B. juncea* var. *tumida* is the raw material of Fuling mustard, which is an important economic vegetable in China and famous for its special flavour and nutritional value ^[17]^. *P. brassicae* leads to the formation of clubroot in tuber mustard, which is one of the most serious stresses and results in the inhibition in plant growth and stem swollen, and leading to limitation in yield ^[18, 31]^. Clade A PP2C family genes play important roles in the regulation of plant development and tolerance to stress, however, the identity and expression patterns of *B. juncea* var. *tumida* clade A *PP2C* genes are unknown. This study is the first identification of clade A *BjuPP2Cs* in *B. juncea* var. *tumida* and adds a new layer to the function of clade A *BjuPP2Cs*.

In this study, twenty clade A *BjuPP2Cs* genes were identified and located in 13 of 18 chromosomes based on the nine clade A AtPP2Cs sequences in *Brassica Database* (Figure 1). *ABI1* had five homologous genes located in A01, A03, B05, A08 and B03; *ABI2* had two homologous genes located in B02 and A10; *AHG1* had two homologous genes located in A03 and B08; *AHG3* had three homologous genes located in A05, B07 and B01; *HAB1* and *HAB2* both had one homologous gene located in the same chromosome B03; *HAI1* had two homologous genes located in A10 and B02; *HAI2* had three homologous genes located in A09, B06 and A10 (Figure 1, Table 1). The comparable homologous gene numbers in A sub-genome and B sub-genome suggested tuber mustard genome experienced co-linearity gene duplication. However, the homologous genes of *HAB1*, *HAB2* and *HAI3* were not duplicated or lost in tuber mustard, indicating that these genes existed a functional redundancy or divarication during the evolutionary process. The losses of genes during the genome duplication events also frequently exist in tuber mustard and other *Brassica* species, such as *BjuPYLs* family genes in tuber mustard ^[32]^ and chitinase family genes in *B. rapa* ^[33]^. The promoters of clade A *BjuPP2Cs* family genes contained various of cis-acting elements, such as ABRE, GT1GMSCAM4, ARFAT and MYB1AT, which regulated plant response to abiotic and biotic stresses and hormone, indicating that the expression level of clade A *BjuPP2Cs* were regulated by hormone, abiotic and biotic stresses (Figure 4). Further we analyzed the expression pattern of clade A *BjuPP2Cs* under pathogen *P. brassicae*. The result showed that *BjuABI1-5*, *BjuAHG1-1*, *BjuAHG3-2*, *BjuAHG3-3*, *BjuHAB1*, *BjuHAB2*, *BjuHAI1-1*, *BjuHAI2-1*, *BjuHAI2-3* and *BjuHAI3* were highly induced by *P. brassicae*, suggesting that these genes regulated tuber mustard response to *P. brassicae* to mediate the formation of clubroot (Figure 7). The stem swollen of tuber mustard were divided into five stages, named D1 (the stems of 1-month-old seedlings, six leaf stage), D2 (the stems of 2-month-old seedlings, primary stage of stem swelling), D3 (the stems of 3-month-old seedlings, early stage of stem swelling), D4 (the stems of 4.5-month-old seedlings, fast-growing stage of stem swelling) and D5 (the stems of 5-month-old seedlings, last stage of stem swelling). In this study, we found that *BjuABI1-1*, *BjuAHG1-1*, *BjuHAI2-2*, *BjuHAI2-3* and *BjuHAI3* were inhibited in D2 to D4, in contrast, *BjuABI1-3*, *BjuABI2-1*, *BjuHAB1* and *BjuHAI1-1* were induced in D2 to D4 (Figure 8). The result suggested that clade A *BjuPP2Cs* might play roles in stem swelling of tuber mustard.

Gene duplication contained three evolutionary fates: subfunctionalization, neofunctionalization, or nonfunctionalization ^[14, 34]^. In our study, each of clade A *BjuPP2Cs* contained 1-5 copies with different gene structures and expression patterns. For example, *BjuABI1-1*, *BjuABI1-2* and *BjuABI1-3* contained 4 exons which is in accordance with *AtABI1*, whereas, *BjuABI1-4* and *BjuABI1-5* only contained 3 exons (Figure 2). The Serine/threonine phosphatases domain was all existed in BjuABI1s proteins (Figure 3). However, the expression patterns of *BjuABI1s* were significantly different, such as all of *BjuABI1s* except *BjuABI1-5* were induced by ABA and only *BjuABI1-2*, *BjuABI1-3* and *BjuABI1-4* were induced by low temperature (Figure 6).

Gene expression analysis showed differences in clade A BjuPP2C genes expression pattern in tuber mustard. Our results indicated that clade A PP2C-based stem swollen regulation and *P. brassicae* response in tuber mustard has evolved distinctly. Different expression patterns of clade A *BjuPP2C* genes to NaCl stress, low temperature and ABA treatment indicated low conservation of gene expression patterns and functional divergence between *B. juncea* var. *tumida* and *A. thaliana* homologous genes.

## Conclusions

Taken together, 20 clade A *BjuPP2Cs* were identified in tuber mustard and the transcript levels of clade A *BjuPP2Cs* under pathogen treatment, abiotic stresses treatment and different development stages were analyzed. The results indicated that the clade A *BjuPP2Cs* might play crucial role in regulating the formation of clubroot, swollen stem and response to abiotic stresses.

## Abbreviations

ABA: Abscisic acid
PP2C: Protein Phosphatase 2C
ABI1: ABA INSENSITIVE 1
ABI2: ABA INSENSITIVE 2
AHG1: ABA-HYPERSENSITIVE GERMINATION 1
AHG3: ABA-HYPERSENSITIVE GERMINATION 3
HAB1: HYPERSENSITIVE TO ABA1
HAB2: HOMOLOGY TO ABI2
HAI1: HIGHLY ABA-INDUCED PP2C GENE 1
HAI2: HIGHLY ABA-INDUCED PP2C GENE 2
HAI3: HIGHLY ABA-INDUCED PP2C GENE 3
PPs: Protein Phosphatases
PPP: Phosphor-Protein Phosphatase
PPM: Phosphoprotein Metallophosphatases
SnRK2s: Sucrose Nonfermenting-1-related Protein Kinase Class 2

## Author contribution statement

CL and CC conceived the project and designed the experiments. YZ, LY, LC, LW, ZC, GC and RS performed the experiments and analyzed the data. CL and CC prepared the manuscript. CC and CL revised the manuscript. All authors read and approved the final manuscript.

## Declarations

### Data availability statements

All data generated or analysed during this study are included in this published article and its supplementary information files.

---

**Table S1** The primers used in this study.

**Fig. S1** Multiple alignment of conserved motifs of the clade A PP2C family proteins. The multiple alignment of conserved motifs was constructed using CLUSTALW. The clade A PP2C family proteins showed 11 conserved sequence motifs, named motif 1 to motif 11 along the top (blue font diaplay). Residues conserved of clade A PP2Cs highlighted in red (totally conservation) and yellow (high conservation).

